# Exploring the alternative conformation of a known protein structure based on contact map prediction

**DOI:** 10.1101/2022.06.07.495232

**Authors:** Jiaxuan Li, Lei Wang, Zefeng Zhu, Chen Song

## Abstract

The rapid development of deep learning-based methods has considerably advanced the field of protein structure prediction. The accuracy of predicting the 3D structures of simple proteins is comparable to that of experimentally determined structures, providing broad possibilities for structure-based biological studies. Another critical question is whether and how multistate structures can be predicted from a given protein sequence. In this study, analysis of multiple two-state proteins demonstrated that deep learning-based contact map predictions contain structural information on both states, which suggests that it is probably appropriate to change the target of deep learningbased protein structure prediction from one specific structure to multiple likely structures. Furthermore, by combining deep learning- and physics-based computational methods, we developed a protocol for exploring alternative conformations from a known structure of a given protein, by which we successfully approached the holo-state conformation of a leucine-binding protein from its apo-state structure.

## Introduction

Protein structure prediction has been a fundamental topic in computational biology for decades.^1–3^ Numerous innovative and reliable methods were developed in recent years, specifically deep learning-based methods, which have greatly advanced this field.^4–10^ Particularly, *AlphaFold2* demonstrated a breakthrough improvement in the accuracy of *de novo* structure prediction in the latest 14*^th^* Critical Assessment of Techniques for Protein Structure Prediction (CASP14).^11,12^ The use of large sequence databases has benefited from the development of bioinformatics; the end-to-end deep neural network outperformed other models in 3D protein structure prediction, reaching high accuracies comparable to those of many experimentally obtained structures. Additionally, new approaches have combined deep learning with traditional computational methods, such as template-based modeling, physics-based optimization, and molecular dynamics (MD) simulations, to achieve improved performance.^13–17^

Currently, most if not all deep-learning-based methods focus on the prediction of one structure in a specific state rather than likely structures of multiple states, while multiple states are essential for most protein functions.^2,18^ For example, many crucial proteins such as enzymes,^19^ G protein-coupled receptors (GPCRs),^20^ ion channels,^21^ and transporters^22^ undergo subtle or significant conformational changes from one stable state to another to exert their functions. Therefore, it is important to study the multiple structures of proteins and the dynamic transition between them to fully understand their functions, for which the ability to predict the protein structures of alternative states is essential. There have been attempts of using *AlphaFold2* to predict alternative structures of proteins. del Alamo et al. found that using shallow multiple sequence alignment (MSA) as input can lead *AlphaFold2* to sample intermediate-like conformations of transporters and receptors, particularly GPCRs.^23^ However, there exist the possibilities that shallow MSAs were not informative enough to generate a converged structure, as pointed out by Heo et al.^24^ Heo and Feig found that the default protocol of *AlphaFold2* shows a strong preference for generating inactive state structures of GPCRs, but one can predict accurate active structures by using state-annotated GPCR structure databases without using MSA.^24^ This is an effective strat-egy to generate activated structures of GPCRs. Nevertheless, it remains elusive whether deep learning actually captured structures of both states or only the dominant one in the structural databases. In addition, the above tests were carried out mostly for GPCRs, whose activation involves relatively small conformational changes. Therefore, further and more extensive validations are required to examine whether deep learning can generally capture the structural information of multiple protein states that can be extracted for a more comprehensive understanding of protein dynamics. From the perspective of physics, a complete understanding of the stable conformational states and transitions among them requires a multidimensional free energy landscape,^25^ which is extremely difficult to obtain. Although experimental and computational methods have advanced in recent years and the number of protein structures in the Protein Data Bank (PDB) has increased markedly,^26–30^ studying large-scale protein conformation transitions remains experimentally and computationally challenging.

Interestingly, direct coupling analysis (DCA) of MSA data can reveal crucial long-range contacts for proteins and protein-protein interactions, which can also be used to explore the likely alternative conformations of proteins and allosteric regulations.^31–34^ Thus, prediction based on DCA may contain conformational information on multiple likely states. Based on pioneering DCA studies, deep learning-based prediction of protein contact maps (CMs) has been improved and widely used for 3D protein structure predictions in the last few years,^6,35,36^ but the biological importance of CM prediction is not as immediately evident as that of DCA. Notably, Iyer et al. found that the difference of CMs from PDB structures can infer the conformational flexibility of proteins,^37^ and Feng et al. developed a method to predict the alternative conformations of proteins by using contacts clustering and Confold2,^38^ strongly suggesting that CM prediction may contain multistate structural information. However, neither the aforementioned *AlphaFold-based* nor CM-based studies excluded the homologous proteins from the training dataset. Therefore, whether and how one can extract residue contact information on multiple likely states from a *de novo* CM prediction by deep learning is still an interesting and promising question.

In this study, by analyzing the predicted CMs and known structures of representative proteins, we demonstrate that it is possible to extract structural information on multiple states from the deep learning predicted CMs. As CM prediction can be viewed as an intermediate step for 3D structure prediction, we think the message is applicable to protein structure prediction as well, i.e., deep-learning-based structure predictions contain information of multiple likely states. In addition, by combining deep learning predictions with physics-based computational methods, we propose a straightforward approach for exploring alternative protein conformations from a known structure guided using CM analysis (Fig. 1). This study will facilitate the prediction of multistate structures based on deep learning methods, as well as accelerate large-scale conformation sampling in MD simulations.

**Figure 1:**
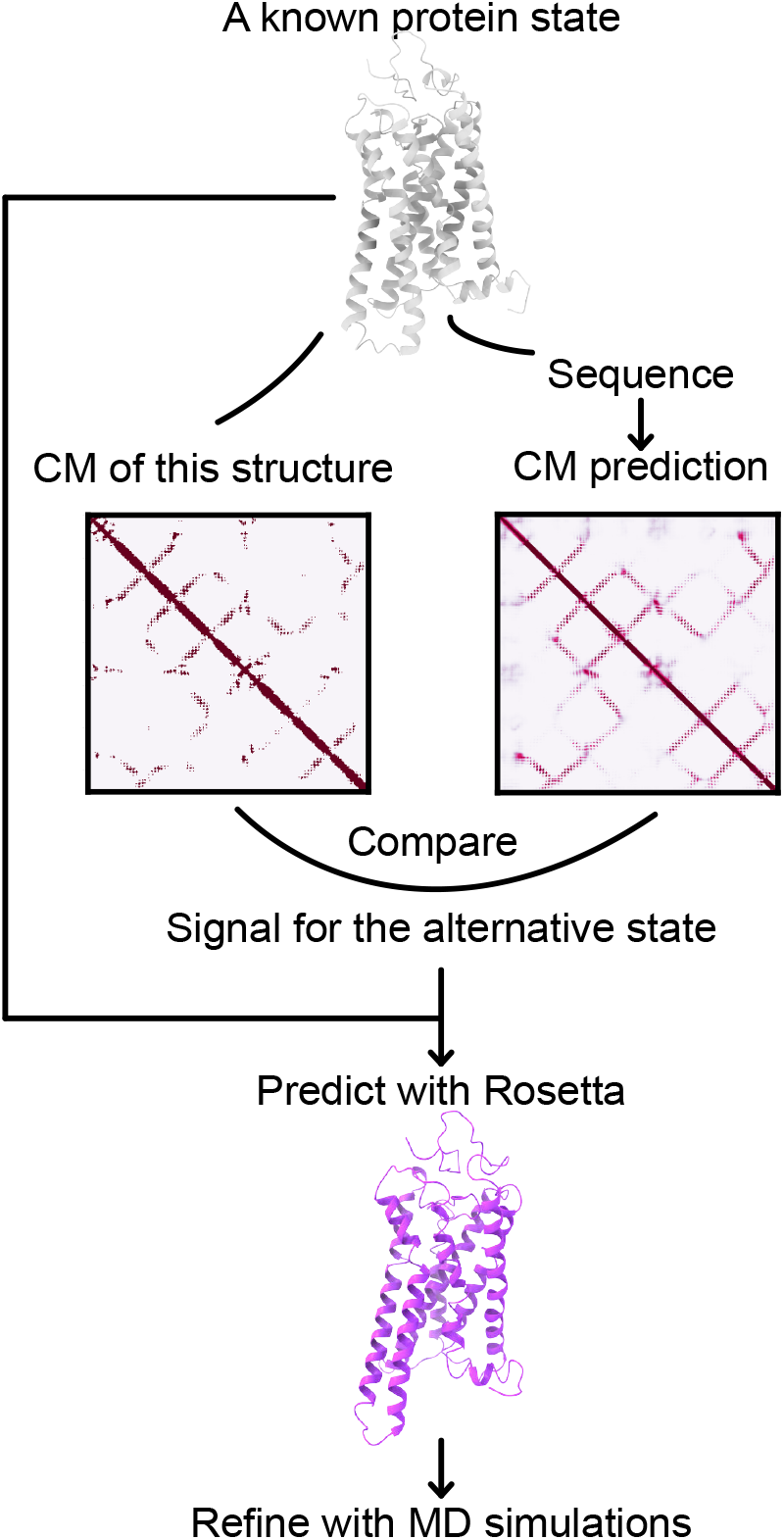
Workflow of our prediction method. We started with a known protein state structure and used the information extracted from its true contact map and predicted contact map to explore the alternative state structure.

## Results

### Predicted contact maps contain structural information on both states for two-state proteins

Two residues are defined as being in direct contact when the Euclidean distance between the *C_β_* atoms (*C_α_* for glycine) is shorter than 8 Å, and the Boolean matrix showing the contact status of all residue pairs in a protein is known as a CM.^39^ Starting from three representative cases, we examined whether protein CMs predicted from sequences based on deep learning and MSA can capture the structural information of both states for two-state proteins. We selected three two-state proteins for CM and structural analyses: rhodopsin, 70-kDa heat shock protein (Hsp70), and leucine-binding protein (LBP). The structures of both states are known for these three proteins and the transitions between their two states involve various degrees of conformational changes, from relatively minor helices rearrangements of rhodopsin to global domain rearrangements of Hsp70.

Our analysis showed that the predicted contact maps indeed contain structural information on multiple states. We calculated the contact maps based on two known structures of both states, which are denoted as TCM1 (true contact map 1) and TCM2. We also used a deep-learning-based method to predict the contact map of the same protein from its sequence (PCM), and then compared the three CMs to check whether the PCMs contain true contacts of both TCM1 and TCM2. The results are summarized in Table 1. As can be seen, when only considering the predictions of “high confidence” (over 0.9), the PCMs of the three proteins not only have the true contacts existing in both states (column 1), but also contain the true contacts that exist exclusively in either state (columns 2 and 3).

**Table 1:** Comparison of predicted contact maps (PCMs) with true contact maps (TCMs) of the two states for the three representative proteins. A three-digit code is used to describe the contact status in the three CMs: the first digit indicates the contact status in TCM1, the second TCM2, and the third PCM. For example, “101” means that a contact exists in the first state structure and the predicted CM, but not in the second state. The values show the number of contacts in the specific status.

|  | Contact status in TCM1, TCM2 and PCM |  |  |  |  |  |  |
| --- | --- | --- | --- | --- | --- | --- | --- |
|  | **111** | **101** | **011** | **001** | **110** | **100** | **010** |
| Rhodopsin | 117 | 17 | 18 | 43 | 263 | 180 | 83 |
| Hsp70 | 756 | 53 | 92 | 149 | 285 | 183 | 274 |
| LBP | 445 | 7 | 16 | 62 | 356 | 42 | 85 |

It is worth mentioning that, the training dataset for our CM predictor did not contain any homologous proteins with a sequence identity over 25% with respect to the target proteins, so the CM predictions can be viewed as *de novo*. Although there are still many false positive (column 4) and false negative (column 5-7) predictions as shown in Table 1, the data in the first three columns suggest that the PCMs can probably be used to construct the structures of both states, if the false positives can be effectively excluded. Removing the false positives from PCMs is a very challenging problem if no other structural information is available. Here we consider an easier scenario, in which the structure of the first state is already known, and we ask the question whether the structure of the second state can be explored based on the known structure and CM analysis. This is a very common scenario, as a protein structure can be readily predicted by deep learning methods nowadays, or even solved experimentally, if it is not already available in the PDB.

### Extracting structural information on the alternative conformation based on a known structure and contact map analysis

By analyzing the above three cases, we established a strict and effective selection criteria to remove the false positives and extract the contact information of the alternative state structure (Materials and methods). In the first case, we analyzed the structures and CMs of rhodopsin, which undergoes a subtle conformational change between its inactive and active states on a few transmembrane helices, mostly the intracellular part of helices 5 and 6 (Fig. 2A). ^40^ Hypothetically, the inactive structure but not the active structure was available. As shown in Fig. 1, we calculated the TCM of the known inactive structure (STR1) and the PCM using the rhodopsin sequence. By subtracting the TCM from the PCM, we obtained the difference contact map (DCM) and identified 61 additional pairs of in-contact residues with prediction confidence of over 0.9 (Fig. 2B), among which 23 pairs were separated by a distance of over 10 Å in STR1 (Table S1). Furthermore, we found 11 pairs of residues that were associated with the conformational change of activation according to our CM analysis criteria, seven of which were located on helices 5 and 6 (Fig. 2A). This result indicates that the additional contacts in the alternative conformation can be extracted by analyzing one known structure and the PCM of rhodopsin, and it is possible to exclude the false positives based on the known structure.

**Figure 2:**
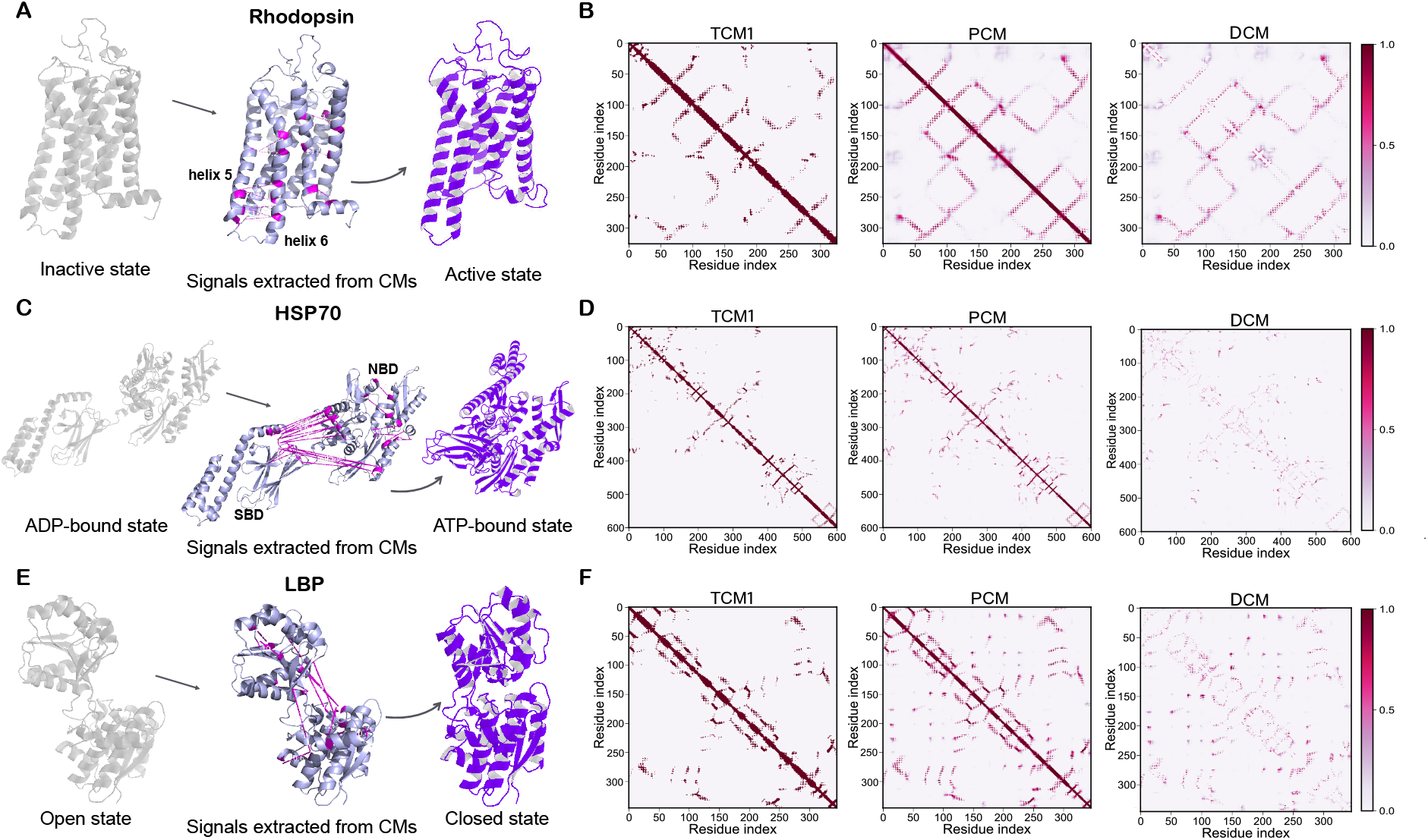
Critical signals associated with conformational changes between two distinct states can be extracted from contact map analyses. (A) Analysis of the transition from the inactive (closed) state (PDB ID: 2i35) to the active (open) state (PDB ID: 6oy9) structures of rhodopsin. (B) Subtracting the true contact map of the known structure (TCM1) from the predicted contact map (PCM) yielded the differences between the two contact maps (DCM), which contains critical signals of likely structural differences when the known structure transitions to the alternative conformation. The selected signals are indicated in (A) with pink lines. (C-D) Similar to (A-B), but for the heat shock protein (HSP70), whose ADP-bound state (PDB ID: 2kho) and ATP-bound state (PDB ID: 4b9q) structures are shown. (E-F) Similar to (A-B), but for leucine-binding protein, the open state (PDB ID: 1usg) and closed state (PDB ID: 1usi) structures are shown.

The second case was heat shock protein (Hsp70), which undergoes large and global con-formational changes when it transitions between the ADP-bound and ATP-bound states (Fig. 2C).^41^ The most evident change was the distance between the nucleotide-binding domain (NBD) and substrate-binding domain (SBD). By following our analysis protocol (Fig. 1) and selection criteria (Materials and methods), we found that: 241 new additional contacts were spotted in the DCM and new clusters of predicted contacts appeared on the PCM compared to in the ADP-bound state (TCM1) (Fig. 2D), indicating that these new regional contacts may exist in the alternative, ATP-bound state (Fig. 2C); 78 pairs of residues with distance over 10 Å were found in the DCM; after ruling out the residue pairs with high flexibility and those showing small distance changes, 27 pairs were selected, which are shown by magenta dashed lines in Fig. 2C. Among these likely contacts, 15 pairs showed an original distance of 20-50 Å (Table S2), indicating that these residues comprised the likely regions of large conformational changes, as they may have to move close to each other until within 8 Å (in contact) in the alternative conformation. Interestingly, the interface between the NBD and SBD in the ATP-bound state was highly correlated with these 15 pairs of residues (Fig. 2C). Therefore, the PCM appears to contain structural information on the two distinct states of Hsp70 during the transition, which involves global and large conformational changes.

We further validated our protocol in the third case, LBP, a periplasmic ligand-binding protein that undergoes a sizable conformational change when transitioning from the apo (open) to the holo (closed) state upon binding to leucine or phenylalanine (Fig. 2E).^42^ Compared to the CM of the open structure (TCM1) of LBP, 78 additional contacts with high prediction confidence (over 0.9) were identified on the PCM (Fig. 2F), showing relatively strong signals for the alternative conformation. According to our selection criteria, 15 pairs of residues were predicted to be in contact in the PCM with a distance (*d*_0_) larger than 10 Å in the known structure (Table 2, Fig. S1), and four pairs of residues (bold in Table 2) were selected for further analysis and modeling. These four contacts may represent the most critical new contacts in the alternative conformation. The distance between these residues is expected to change from above 15 Å to lower than 8 Å in the new conformation, according to the PCM and known structure. In addition, these four pairs of residues were present at the interface of two domains, indicating that the two domains should approach each other in the alternative conformation (Fig. 2E).

**Table 2:**
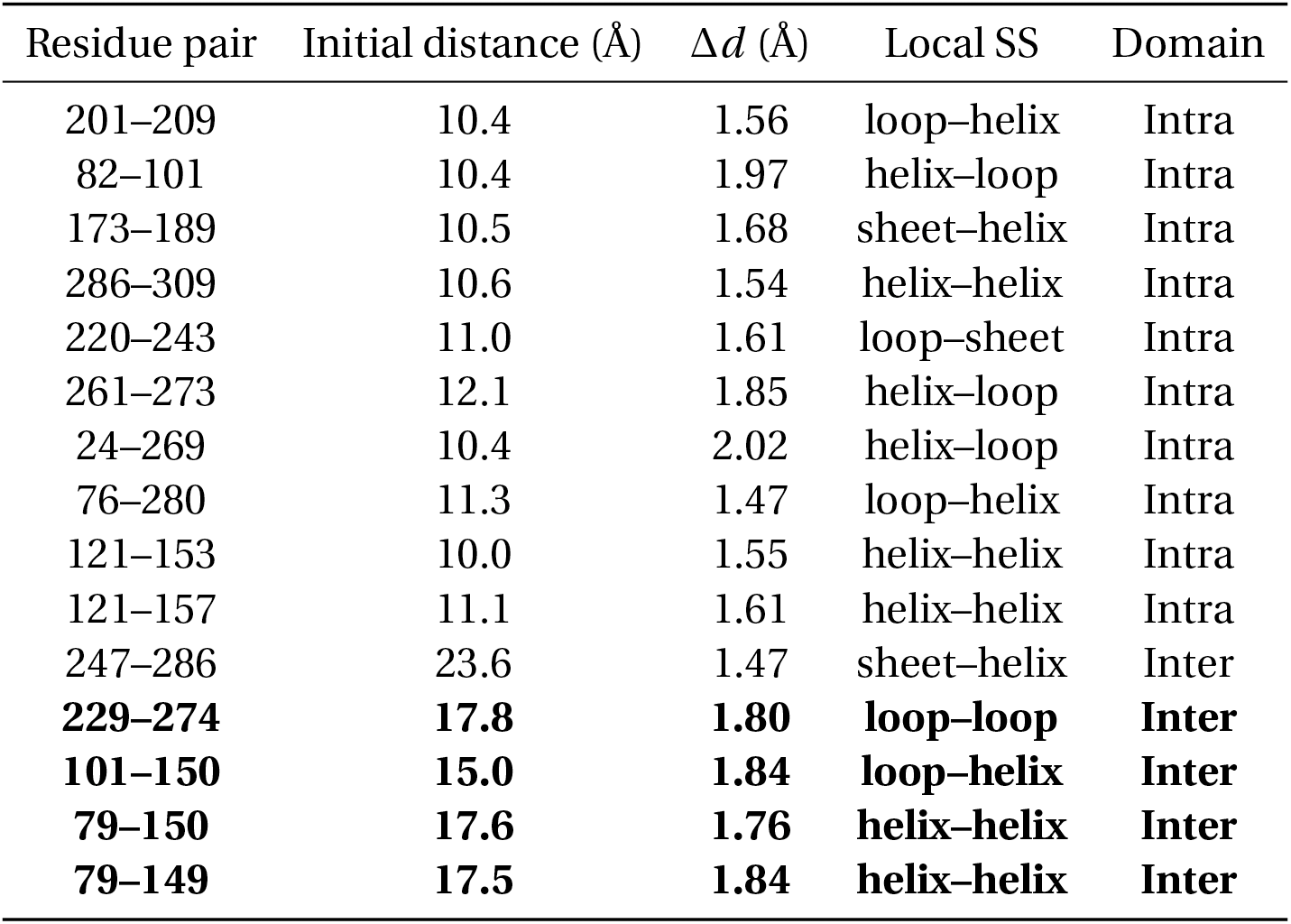
Selection of key residue pairs based on contact map analysis of leucine-binding protein. The final selection is indicated in bold.

| Residue pair | Initial distance (Å) | $\Delta d$ (Å) | Local SS | Domain |
| --- | --- | --- | --- | --- |
| 201–209 | 10.4 | 1.56 | loop–helix | Intra |
| 82–101 | 10.4 | 1.97 | helix–loop | Intra |
| 173–189 | 10.5 | 1.68 | sheet–helix | Intra |
| 286–309 | 10.6 | 1.54 | helix–helix | Intra |
| 220–243 | 11.0 | 1.61 | loop–sheet | Intra |
| 261–273 | 12.1 | 1.85 | helix–loop | Intra |
| 24–269 | 10.4 | 2.02 | helix–loop | Intra |
| 76–280 | 11.3 | 1.47 | loop–helix | Intra |
| 121–153 | 10.0 | 1.55 | helix–helix | Intra |
| 121–157 | 11.1 | 1.61 | helix–helix | Intra |
| 247–286 | 23.6 | 1.47 | sheet–helix | Inter |
| **229–274** | **17.8** | **1.80** | **loop–loop** | **Inter** |
| **101–150** | **15.0** | **1.84** | **loop–helix** | **Inter** |
| **79–150** | **17.6** | **1.76** | **helix–helix** | **Inter** |
| **79–149** | **17.5** | **1.84** | **helix–helix** | **Inter** |

In our selection criteria from the DCM, the potential contact with large distance variation, inter-secondary structures, or inter-domains were strongly preferred. This will exclude some true positives in the DCM, but the advantage is that all the false negatives can be excluded and some key contacts in the alternative conformation can still be identified for further analysis and modeling.

### Prediction of alternative conformation of LBP with Rosetta

We further explored whether the contact information identified above can help predict or model the alternative conformation. We used LBP as a validation system because the structural transition between its two states exhibits significant and straightforward conformational changes (Fig. 2E). Hypothetically, the apo (open) state structure but not the holo (closed) state structure of LBP was available. Based on the above analysis, four critical residue pairs were predicted to be close to each other in the holo state structure (Table 2 and Fig. 3A). Such a conformation exploration problem can be solved using a physics-based optimization method with a powerful tool such as the widely used Rosetta software suite. We constructed an energy-distance function according to the likely contacts of the alternative conformation (Fig. 3B) to guide the selected residue pairs to approach each other to match the above contact signals from the DCM. After adding this energy function to the optimization procedure of Rosetta, a closed-state model of LBP was obtained from the open-state crystal structure (details in Materials and methods).

**Figure 3:**
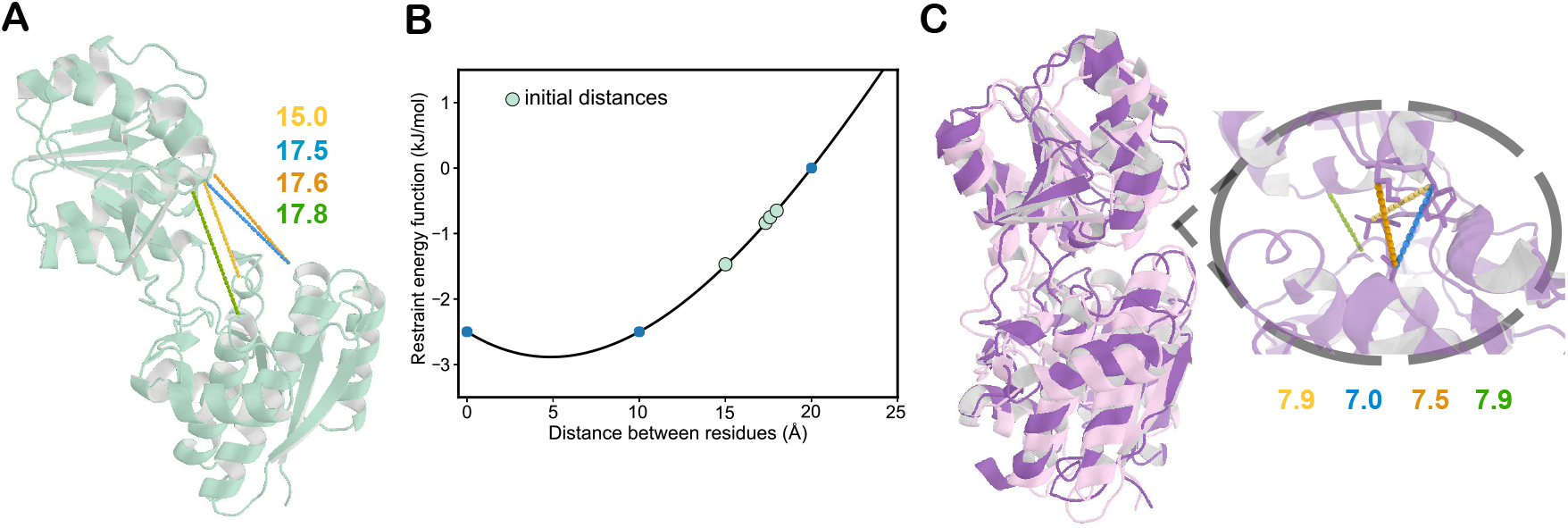
Prediction of the closed state structure of LBP from the open structure. (A) Distances (inÅ) between the four selected pairs of residues in the original open-state structure. (B) Restraint energy function for the four selected residue pairs for the optimization procedure in Rosetta. (C) Predicted closed structure (purple) aligned with the holo-state crystal structure (light pink) with a TM-score of 0.81.

As shown in Fig. 3C, we obtained a closed conformation of LBP from its open state structure. The quantitative change in distances between the four pairs of residues was labeled. Comparisons of the predicted conformation with the crystal structures of the two states, as evaluated using the template modeling score (TM-score) and root-mean-square deviation (RMSD), are listed in Table 3. The predicted structure resembled the holo-state crystal structure (TM-score: 0.81; RMSD: 3.6 Å) while deviating significantly from the initial model — the open-state crystal structure (TM-score: 0.62; RMSD: 6.7 Å), indicating that a rough holo-state model was obtained.

**Table 3:** TM-score and RMSD of the predicted or sampled conformations of leucine-binding protein, with respect to its open and closed structures, respectively.

| Model | TM-score |  | RMSD (Å) |  |
| --- | --- | --- | --- | --- |
|  | Open state<br>(1usg) | Closed state<br>(1usi) | Open state<br>(1usg) | Closed state<br>(1usi) |
| Closed state (predicted) | 0.62 | **0.81** | 6.7 | 3.6 |
| MD1 | 0.60 | **0.87** | 7.6 | 2.0 |
| MD2 | 0.60 | **0.86** | 7.9 | 2.5 |
| MD3 | 0.61 | **0.88** | 7.4 | 2.0 |
| Closed state (1usi) | 0.64 | 1.00 | 7.2 | 0.0 |

### MD refinement of alternative conformation

To further refine the rough model obtained above, atomistic MD simulations were performed. First, we conducted a 500-ns MD simulation using the closed, holo-state crystal structure as the initial configuration. This control simulation showed that the closed state was unstable in the absence of the ligand and tended to quickly evolve back to the open state, with the RMSD of the protein conformation with respect to the holo-state crystal structure reaching above 7 Å (orange line in Fig. 4A). We then performed three independent 500-ns MD simulations to evaluate the stability of our predicted LBP conformation without applying distance restraints. The results were similar to those of the control simulation, and the RMSD with respect to the holo-state structure fluctuated over a large range between 4 and 11 Å (blue lines in Fig. 4A). These results indicate that the predicted conformation was unstable in the absence of ligands or external restraints, which is in accord with expectation.

**Figure 4:**
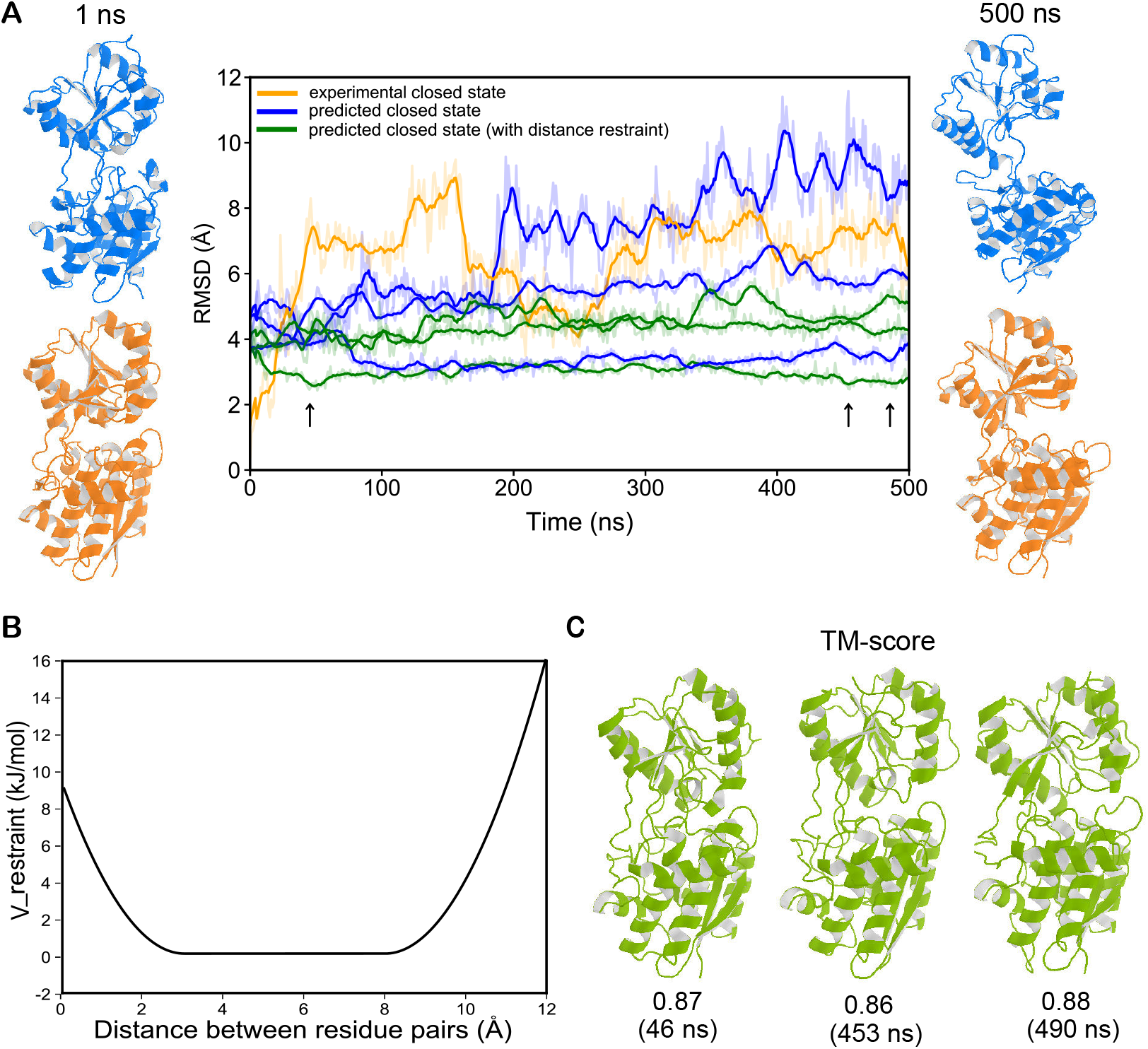
Molecular dynamics (MD) refinement for the predicted structure. (A) The RMSD of 500-ns MD simulations with respect to the closed structure. The simulations started with the experimentally solved closed structure (left orange) and predicted closed structure (left blue), with or without distance restraints. Conformations at 500 ns are also shown for the simulations without distance restraint (right). (B) Distance restraint potential applied to the selected pairs of residues in MD simulations. (C) Three best-sampled conformations in the MD simulations with distance restraint (indicated by arrows in (A)), showing the lowest RMSD values and best TM-scores with respect to the closed structure.

To better refine the predicted closed-state model without having to incorporate the ligand, which would introduce other uncertainties, we applied distance restraints in our MD simulations. We added a piece-wise harmonic function to maintain the distances between the predicted in-contact residue pairs in the MD simulation (Fig. 4B, Materials and methods) to better maintain and refine the conformation of the predicted model from Rosetta. As shown by the green lines in Fig. 4A, there were still deviations from the holo-state structure in the three independent simulations with distance restraints, but the RMSD values fluctuated around 3 to 5 Å (Fig. 4A), indicating that the sampled LBP conformations were more close to the holo state than in the simulations without restraints. Indeed, we obtained multiple conformations that were highly similar to the holo-state crystal structure in our MD trajectories (Fig. 4C and Table 3). In addition, the local conformation of helices within the lower domain was also optimized compared with that of the initial model from Rosetta (Fig. S2). The best-sampled conformation showed a TM-score of 0.88 and an RMSD of 2.0 Å, indicating that conformations highly similar to the holo-state structure had been sampled.

## Discussion

Deep learning-based protein structure prediction has achieved astonishing accuracy for simple proteins in recent years.^8,11^ However, most existing methods focus on predicting a specific structure without considering the existence of multiple-state conformations. Recently, *AlphaFold2*’s prediction performance on different conformations has also drawn much attention. Besides the attempts of two groups to predict the alternative conformations of GPCRs,^23,24^ Saldaño et al. elaborately collected 91 pairs of apo-holo structures and used them to examine whether *AlphaFold2* could capture those two-state conformations.^43^ It was found that *AlphaFold2* failed to yield models resembling different functional conformations for the given protein sequence. Nonetheless, this dataset provides an ideal test set for our method as well. By analyzing those 91 pairs of proteins, we further proved that CMs predicted via deep learning indeed contain structural information on both states in most cases (Fig. 5 and Table S3). The predicted CMs of 84 proteins in the 91-protein dataset contain structural information of both states, indicating that the current deep learning-based predictions work fairly well in capturing multistate structural information, although there is still room for further improvement. The fact that the multistate conformation can be predicted from the CMs probably relies on the exploration of MSA information by machine learningbased approaches. This occurs because diverse stable or metastable states of proteins are selected by evolution, and the evolutionarily important residue contacts can be captured, regardless of whether the interaction stabilizes one or the other states. In fact, similar ideas were proposed in the pioneering work by other groups based on DCA.^31,32,44^ Here we further expand the idea to deep-learning-based CM prediction and provide strong evidence for *de novo* prediction of multistate structural information by using deep learning, in addition to Feng and Shukla’s work that utilized a different strategy.^38^

**Figure 5:**
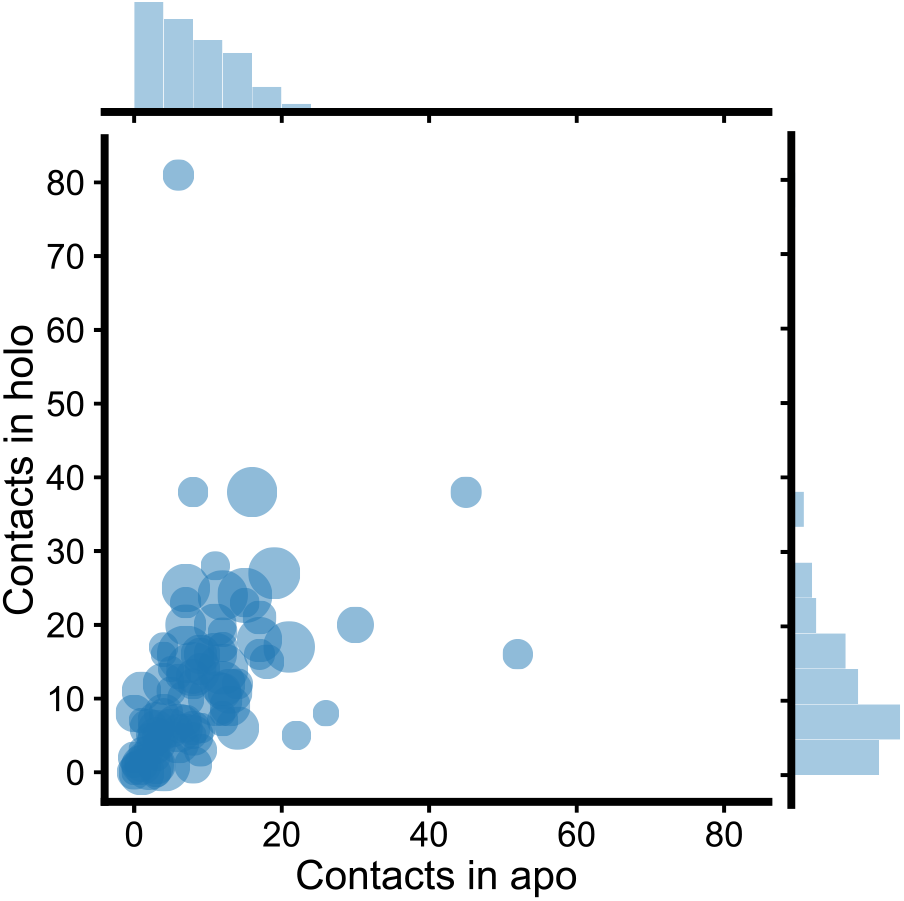
Analysis of contacts in the predicted contact maps (PCMs) for the additional 91 proteins. The number of apo/holo-state specific contacts (“101”/”011” in table S3) were represented on the x/y axis. The size of circles represented proteins’ sequence length. The PCMs of most proteins cover certain amounts of the state-specific contacts of both states.

Although the CM only shows 2D information, they have been extensively used for constructing 3D structures and can be predicted in parallel with 3D structures.^36,45^ Therefore, the conclusion may be applicable for 3D protein structure prediction too; that is, knowledge-based deep-learning prediction may be used for multistate structure prediction. This implies that current deep learning-based protein structure predictions can probably be further improved by adjusting the prediction target from one specific structure to multiple likely structures, which could not only improve the prediction accuracy but also expand the biological significance of protein structure predictions.

Physics-based computer simulations of proteins have been greatly hindered by difficulties related to efficient sampling. Specifically, MD simulations are effective for sampling the conformational space around the initial configuration and can often generate quantitative results consistent with the functional state of the initial structure. However, it remains challenging to simulate the global conformational transition from one state to another, particularly when the transition process faces a high free energy barrier and prior knowledge of the reaction coordinate is lacking. In this regard, our approach provides a potential solution for determining the reaction coordinate or conformation sampling direction by analyzing deep learning predictions. This protocol can be used to sample the alternative conformation of a protein based on its known structure and PCM. In fact, similar methods can be used with other types of predictions, such as torsion angle and accessibility predictions, to drive large-scale conformational transitions; however, deep learning predictions of global structural features would probably be more useful for steering major and global conformational changes. There are still challenges though: due to the absence of the binding ligand and limitation of sampling efficiency, the best sampled conformation was not the most probable conformation in MD simulations in the LBP case. Therefore, more sophisticated refinement protocol need to be developed and utilized in future studies.

The criteria used to extract information from PCMs in this study have proven useful for multiple protein systems, including both local conformational changes within one domain and global changes across domains. However, several parameters in the method depend on the specific properties of the protein being evaluated, such as its size, secondary structure, flexibility, domain composition, and surrounding environment of the protein. Therefore, some prior knowledge of the protein is still necessary for establishing an efficient protocol. Fortunately, most of this information can be obtained by the experimentally resolved structures or deep learning-based predictions now. In addition, the method for constructing the energy-distance function and distance restraint strength can also be further improved to explore alternative conformations. For instance, a predicted distance map, rather than a CM, is useful for constructing a more accurate energy-distance function. It is encouraging to see that, based on the structural information of the known state, most false positives in the PCM can be excluded. Obviously, the protocol can also be used with a reliably predicted structure as the known state, such as a predicted structure by *AlphaFold2* or *RoseTTAFold.* Therefore, in principle, our method and its derivatives can be used in combination with existing 3D structure prediction tools to explore multiple likely protein structures, which can be further combined with MD simulations to depict the free energy landscape of protein dynamics.

A more challenging question is how to deal with proteins with more than two states when analyzing the PCMs and clustering contacts during the 3D structure prediction procedure. Some methods for self-consistency checks may be useful, which require further examination. Meanwhile, previous studies have shown that the development of MD enhanced sampling together with elaborate structure refinement procedures would probably also contribute to the prediction of alternative conformations.^15,16,28,46^ Therefore, the combination of knowledge-based deep learning predictions and physics-based computer simulations is expected to provide new possibilities and more comprehensive pictures in the study of multistate structures and free energy landscapes of proteins.

In summary, this study revealed that multiple-state structural information can be extracted from *de novo* deep learning predictions, which can be further utilized for structural modeling based on physics-based approaches and constructing reaction coordinates for efficient sampling in MD simulations. This method can be improved, but the framework is sufficiently general to be used as a quick test of hypotheses related to protein structure changes and can be used in combination with experimental techniques to study protein dynamics. The method will be useful for providing biophysical insights into the underlying structural and dynamic mechanisms of multistate proteins.

## Materials and methods

### Overall protocol

To determine whether the PCMs contain structural information for multiple states, we evaluated three representative two-state proteins whose structures in both states have been resolved. The overall protocol is as follows.

1. As shown in Fig. 1, starting with one known structure (STR1), we directly calculated its contact map, which was named as “TCM1” (true contact map 1).
2. We used a deep learning-based method to predict the CM from the sequence of the same structure,^4,47^ named as “PCM”.
3. The matrix of the TCM1 was subtracted from PCM, and the resulting contact map was named “DCM” (difference contact map).
4. The CM of the structure of the second state (STR2) was calculated directly from the known structure and named as TCM2 (true contact map 2). Please note that this step is only for the purpose of validation and is not required for prediction or modeling.
5. We examined whether the DCM contained the contact information of TCM2 that did not exist in TCM1, and extracted the contact information for further structure modeling.

After identifying key contact information in the DCM, we applied additional energy-distance restraint functions in the physics-based model-building tools to search for an alternative structure (STR2) from the known structure (STR1):

1. Starting with STR1, we added energy-distance restraint functions to the ‘initialize pose’ module of Rosetta and searched for an alternative structural model (see below for details).
2. With the structure generated using Rosetta, we performed MD simulations with distance restraints to further refine the structure (see below for details).

### Protein structures studied in this work

#### Rhodopsin

GPCRs form a large group of membrane-embedded receptor proteins that are involved in a plethora of diverse processes.^20^ Rhodopsin, a class A GPCR, is the light receptor in rod photoreceptor cells of the retina and plays a central role in phototransduction and rod photoreceptor cell health.^48^ Upon activation, rhodopsin undergoes subtle conformational
changes in a few transmembrane helices. The inactive (PDB ID: 2i35) and active (PDB ID: 6oy9) structures have both been resolved and deposited in the Protein Data Bank (PDB), which were downloaded for analysis.

#### Heat shock protein (Hsp70)

The 70-kDa heat-shock proteins (Hsp70s) are ubiquitous molecular chaperones essential for cellular protein folding and proteostasis.^49,50^ Hsp70 has two functional domains: a nucleotide-binding domain and a substrate-binding domain. The formation of domain interfaces is associated with significant conformational changes upon binding to ATP and polypeptide substrates. The structures of the ADP-bound (PDB ID: 2kho) and ATP-bound states (PDB ID: 4b9q) were downloaded from the PDB.

#### Leucine-binding protein (LBP)

The periplasmic leucine-binding protein is the primary receptor for the leucine transport system in *Escherichia coli.^51^* The crystal structures of both the apo and holo forms have been resolved and have revealed that an ‘open-to-close’ conformational change was associated with ligand binding. The open (PDB ID: 1usg) and closed (PDB ID: 1usi) state structures were downloaded from the PDB.

#### The additional 91 two-state proteins

To make more comprehensive tests, we adopted the 91 two-state proteins curated by a recently published work,^43^ in which the structures of both apo and holo states were available for each of the 91 proteins.

### Protein CM prediction

In recent years, machine learning-based methods for contact map (CM) prediction have advanced significantly and become the mainstream in the field of CM prediction. Xu et al. developed a ResNet-based model for the CM prediction (RaptorX-Contact), which showed breakthrough performances.^4^ Based on this deep model, we built a ResNet-based CM predictor with a new character of proteins, membrane contact probability (MCP), incorporated to better account for the structural feature of membrane proteins and achieved improved performance for both soluble and membrane proteins.^47^

In this study, we updated the training data set and trained new MCP-incorporated contact map predictors with the same hyperparameters as in our previous work.^47^ For the three representative proteins, we removed the redundant sequences in the training sets with a strict criterion so that there were no training proteins with sequence identity > 25% with respect to any of them. For the 91 additional proteins, we also removed their homologous sequences according to the same criterion and re-trained the model. To predict the CM of each protein from its sequence, we ran HHblits 3.0.3^52^ (with an E-value of 0.001 and three iterations) to find its homologous in the Uniclust30 database dated October 2017 and built its multiple sequence alignment (MSA). Then the input features were generated from the MSA and put into the ResNet model for the CM prediction. Please refer to the previous publications for more details.^4,47^

### Analysis of true contact maps and predicted contact maps

For a certain residue pair of a given two-state protein, their contact status in the two TCMs and the PCM can be expressed by a three-digit code. The first digit corresponds to the contact status in TCM1, the second in TCM2, and the third in PCM. “1” means there is a contact and “0” no contact. Therefore, “101” means there is a contact in TCM1 and PCM, but not in TCM2. There are eight different sets to show the contact status of the residue pair in the three CMs, but “000” is not of interest and hence discarded. The number of each status were counted for all the 94 proteins. The analysis results for the three representative and the 91 additional two-state proteins are shown in Table 1 and Table S3, respectively. Noted that we only include residue pairs with sequence spacing ≥ 6 for counting, and only considered those contacts in the PCM with a prediction confidence of > 0.9.

### Extracting information on alternative conformations from CM analysis

To compare the CM of the known structure (TCM1) with the PCM and extract useful information, the following procedure was performed.

1. The PCM was subtracted by TCM1 to yield the DCM, in which only the residue pairs
predicted to be in contact (distance between *C_β_* atoms < 8 Å) with a confidence *p* > 0.9 in the PCM were considered. The residue pairs were categorized into three groups according to their spacing in the protein sequence, named as short-, medium-, or long-range, if the sequence distance fell into [6, 11], [12, 23], and ≥ 24, respectively. The medium- and long-range contacts were of primary interest because they may help identify larger conformational changes. Therefore, the predicted medium- and long-range contacts with high confidence that did not exist in TCM1 were further analyzed.
2. The above extracted contact information may indicate which residue pairs have direct contacts in the alternative conformation, and only residue pairs with distances over 10 Å in the STR1 (*d_0_*) were selected for further operation. The cut-off distance in the CM definition is 8 Å, and thus residue pairs with distances lower than 10 Å (*d*_0_ < 10) may vary little and have a weak impact on the global conformational change of the protein when searching for an alternative conformation, and thus were discarded before further analysis.
3. The flexibility of the known structure (STR1) was also considered. The b-factor values (B) in the PDB file of STR1 reflect the fluctuation of atoms in their average positions and provide valuable information on protein flexibility. The root-mean-square fluctuation (RMSF) was calculated using Equation 1, in which 〈*B*〉 represented the average value of all atoms in one residue. And the closest residue-residue distance *d_min_* was calculated using Equation 2 to account for the flexibility of each residue pair.

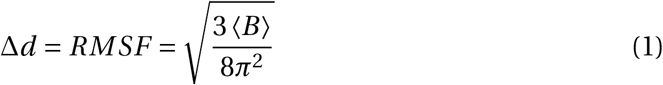

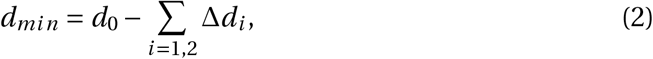

where *d*_0_ represents the distance between the residue pairs in the known protein structure (STR1). The residue pairs with *d_min_*< 10 (Å) was discarded for further analysis as well. This step assists in screening out noise caused by protein flexibility, which is particularly helpful for the analysis of small conformational changes.
4. The rigidity of local secondary structures was taken into account. Different types of secondary structures show varying degrees of conformational flexibility,^53^ and *α*-helices and *β*-sheets are more rigid than loops; therefore, the residue pairs within one *α*-helix or *β*-sheet that are predicted to be in contact with the DCMs were excluded from further analysis; the residue pairs separated by stable, local structures were also discarded to avoid disruption within a structural domain.

Following the above procedure, false positive and ambiguous signals were screened out.

### Model building and optimization

#### Initial modeling with Rosetta

Starting with the STR1 of LBP (PDB ID: 1usg, resolution: 1.53 Å), we used Rosetta to build an approximate model of the alternative conformation with restraints derived from the above DCM signals. Three functional modules of PyRosetta,^54^ pose initialization, minimization, and full-atom refinement, were employed in this process.

First, we initialized the pose of our input structure (STR1) with the switch type as “centroid”. Specifically, a mutation of residue GLY to ALA was adopted in our protocol to account for the lack of *C_β_* atoms in GLY. This made the protein suitable for adding a distance-energy function in the following steps, as the distance between residues was calculated from the positions of *C_β_* atoms.

For the minimization stage, we designed a piece-wise energy-distance function to represent the restraints derived from the DCM signals, as a previous study showed that such a function is suitable for modeling protein structures.^6^ The restraint energy-distance function was as follows:

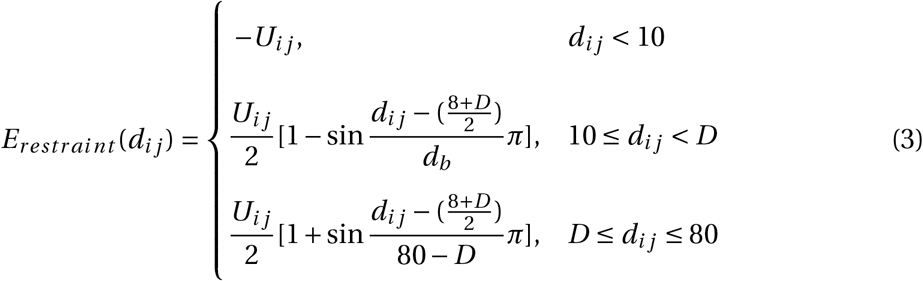

where

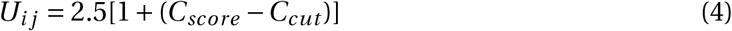

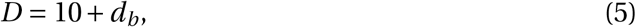

In Equations 3–5, *U_ij_* represents the intensity of the functions, which is related to the precision of the PCM (*C_score_*) and cut-off confidence value (*C_cut_*) chosen prior to analysis (typically set to 0.9). D represents the boundary of the distance interval, as expressed in Equation 3. *d_ij_* represents the distance between the *C_β_* atoms of the indexed residue pairs, and *d_b_* represents the width of the energy barrier, which is determined from the length of the protein sequence: *d_b_* = 10 Å for protein lengths over 250 amino acids, 8 Å for protein lengths between 200 and 250 amino acids, and 6 Å for protein lengths below 200 amino acids.

After the restraint energy score from the DCM was calculated using Equation 3, the discrete scores were converted into a smooth energy potential using the *spline* function in Rosetta and used as restraints to guide energy minimization and structure modeling.

We then used quasi-Newton-based energy minimization function^55^ of PyRosetta (MinMover) to lower the energy of the protein backbones and build coarse-grained models. At this stage, protein structures were represented using a centroid model, in which the side chains were simplified into single artificial atoms (centroids), whereas the backbones remained atomistic. Optimization was performed based on the L-BFGS algorithm (lbfgs_armijo_nonmonoto with a maximum of 1,000 iterations, and the convergence cut-off was set to 0.0001. In addition to the *spline* function mentioned previously, several Rosetta energy forms were used, including centroid backbone hydrogen bonding (cen_hb), ramachandran (rama), omega, and steric repulsion van der Waals forces. The weights of cen_hb, rama, omega, and van der Waals were 5, 1, 0.5, and 3, respectively. The orientation distributions were determined according to the protocol of trRosetta to set *dihedral* and *angle* restraints with equal weights. Detailed potential instructions for *θ, ω* (*dihedral*), and *ϕ* (*angle*) can be found in a previous publication.^45^

Finally, we used the relaxation function (FastRelax) to generate the predicted full-atom model. The top 50 models with the lowest energies built using MinMover were used for FastRelax. During relaxation, both distance restraints (the *spline* function) and orientation restraints explained in MinMover were added to the side chains. We used the score function *ref2015* from Rosetta^56^ to obtain physically plausible conformations. A maximum iteration of 200 was sufficient for FastRelax with a flexible MoveMap to generate the final model of the alternative structure.

#### Conformation sampling with MD simulations

Using the approximate model obtained above, we further performed MD simulations to evaluate the stability of the predicted LBP conformations and applied step-wise refinements to obtain better sampling for the alternative conformation (holo, closed state). All simulations were performed using GROMACS 2018.4^57^ and CHARMM36m force field.^58^ Each protein system was solvated in a water box of 8.9 nm × 8.9 nm × 8.9 nm with periodic boundary conditions, and Na^+^ was added to the solvent to neutralize the simulation system.

We performed two rounds of equilibration before the production simulations. First, we used the steepest descent algorithm for 50,000 steps to minimize the system energy. Next, the system underwent a 100-ps equilibration to reach 310 K in the canonical ensemble (NVT) and 500-ps equilibration to reach a pressure of 1.0 bar in the NPT ensemble using the Berendsen coupling method.^59^ Position restraints were applied to the protein backbone (force constant 1000 kJ mol^−1^nm^−2^). The time step of the equilibration was 2 fs. The particle-mesh Ewald method^60^ was used to calculate long-range electrostatic interactions. The van der Waals interactions were smoothly switched off from 1.0 to 1.2 nm. An elastic network was used to maintain the global conformation of proteins during equilibration.

We then used virtual sites^61^ to increase the time step from 2 to 4 fs for the sake of simulation efficiency in the production simulations. Again, energy minimization followed by a 100-ps NVT equilibration and 500-ps NPT ensemble equilibration were conducted before the final production simulations.

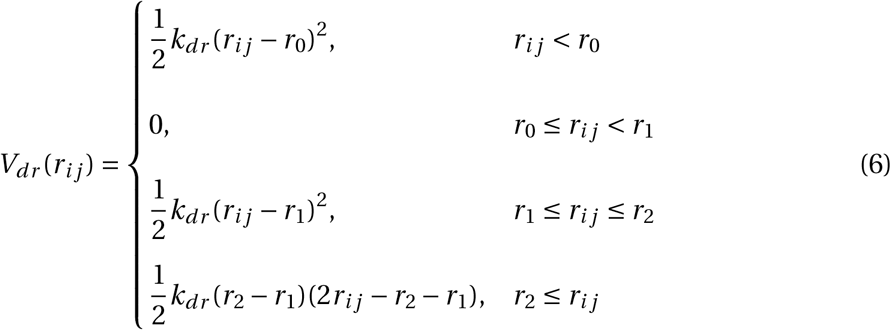

To achieve the aforementioned potential contacts in the alternative conformation (four residue pairs for LBP shown in Table 2 and Fig. 3B), in some of the production MD simulations, we added distance restraints using a piecewise harmonic function as described in Equation 6, where *k_dr_* was set to 200 kJ mol^−1^ nm^−2^; *r*_0_, *r*_1_, and *r*_2_ were set to 0.3, 0.8, and 1.2 nm, respectively. Please refer to the *Distance restraints* section in the GROMACS documentation for more details. Meanwhile, all the other restraints were removed. The systems were simulated for 500 ns multiple times and analyzed using VMD^62^ and PyMOL (https://pymol.org/).

## Acknowledgement

We thank Prof. Bert de Groot for helpful discussions on CONCOORD. This work was supported by grants from the Ministry of Science and Technology of China (2021YFE0108100) and National Natural Science Foundation of China (21873006 and 32071251). Part of the MD simulations were performed on the computing platform of the Center for Life Sciences at Peking University.

## Author contributions

C.S. designed and supervised the project. J.L. conducted CM analysis, structural modeling, and MD simulations. L.W. trained the CM predictor and made CM predictions. Z.Z. analyzed the CMs of the additional 91-protein dataset. All the authors participated in the writing of the manuscript.

