## Supplementary Tables and Figures for "Exploring the alternative conformation of a known protein structure based on contact map prediction"

**Table S1:** The 23 pairs of residues with distance over 10 Å that are likely in contact (prediction confidence over 0.9) in the active state but not in the inactive state of rhodopsin. The selected residue pairs according to our criteria are indicated in bold, which are shown by pink lines in Fig. 2A.

| | **Residue pairs** | **Initial distance (Å)** |
|---|---|---|
| **Short-range** | 138–145 | 13.2 |
| | **308–317** | **11.4** |
| **Medium-range** | **227–251** | **10.9** |
| | **230–247** | **10.3** |
| | **230–250** | **12.8** |
| | **234–247** | **10.3** |
| **Long-range** | **44–292** | **11.3** |
| | **45–94** | **10.6** |
| | 47–90 | 10.4 |
| | **51–298** | **11.5** |
| | 79–161 | 11.3 |
| | 82–165 | 11.4 |
| | 89–120 | 11.5 |
| | 110–185 | 10.3 |
| | 120–300 | 11.1 |
| | 123–299 | 10.5 |
| | 123–303 | 13.4 |
| | 124–303 | 11.1 |
| | 127–303 | 12.7 |
| | 131–313 | 15.0 |
| | **215–261** | **10.2** |
| | **215–265** | **11.4** |
| | **227–254** | **13.3** |

**Table S2:** The 15 residue pairs that may be associated with the significant conformational change of Hsp70 from the ADP-bound to the ATP-bound state from our analysis.

| Residue pairs | Initial distance (Å) |
|---|---|
| 148–506 | 35.7 |
| 149–506 | 39.1 |
| 152–506 | 33.4 |
| 152–507 | 31.3 |
| 156–507 | 32.2 |
| 156–512 | 28.6 |
| 159–512 | 24.1 |
| 160–512 | 28.8 |
| 160–515 | 34.0 |
| 160–516 | 31.4 |
| 217–392 | 27.0 |
| 326–414 | 49.5 |
| 326–415 | 45.6 |
| 327–414 | 48.2 |
| 327–415 | 44.6 |

**Table S3:** Contact map (CM) analysis for the additional 91 protein, sorted by protein sequence length (from long to short). Similar to Table 1, a three-digit code is used to describe the contact status in the three CMs: the first digit indicates the contact status in TCM1, the second TCM2, and the third PCM. For example, "101" means that a contact exists in the first state structure and the predicted CM, but not in the second state. The values show the number of contacts in the specific status.

| Protein Information | 111 | 101 | 011 | 001 | 110 | 100 | 010 |
|---|---|---|---|---|---|---|---|
| 1L0W_B–1G51_A | 441 | 7 | 16 | 112 | 564 | 31 | 124 |
| 7AZP_A–4PJ1_A | 265 | 15 | 24 | 53 | 565 | 135 | 227 |
| 1RKM_A–1B6H_A | 460 | 11 | 10 | 119 | 565 | 55 | 102 |
| 1A6D_A–1A6E_A | 436 | 8 | 14 | 74 | 517 | 53 | 72 |
| 1Y3Q_A–1Y3N_A | 283 | 4 | 1 | 93 | 558 | 50 | 71 |
| 1W0J_E–1E1R_F | 380 | 19 | 27 | 60 | 476 | 89 | 94 |
| 2BNH_A–1DFJ_I | 586 | 21 | 17 | 67 | 420 | 56 | 80 |
| 1E5L_A–1E5Q_A | 416 | 12 | 14 | 62 | 471 | 52 | 74 |
| 1HOO_B–1CG0_A | 446 | 12 | 24 | 123 | 409 | 53 | 127 |
| 1RF5_A–1RF4_A | 503 | 16 | 38 | 159 | 450 | 43 | 117 |
| 1K6W_A–1K70_A | 342 | 2 | 1 | 125 | 568 | 19 | 29 |
| 1EVK_A–1EVL_A | 262 | 6 | 6 | 91 | 447 | 47 | 66 |
| 1K5H_A–1Q0Q_A | 364 | 7 | 25 | 90 | 351 | 89 | 103 |
| 1JEJ_A–1JG6_A | 18 | 1 | 0 | 9 | 615 | 29 | 60 |
| 1Z15_A–1Z17_A | 411 | 12 | 12 | 76 | 283 | 21 | 96 |
| 6OY9_B–1A0R_B | 369 | 17 | 18 | 26 | 266 | 65 | 129 |
| 1VR6_A–1RZM_A | 379 | 13 | 11 | 61 | 311 | 15 | 69 |
| 1ZA1_A–1Q95_A | 390 | 14 | 6 | 107 | 213 | 48 | 62 |
| 1RKA_A–1GQT_A | 419 | 11 | 16 | 42 | 275 | 26 | 29 |
| 4LP5_A–4P2Y_A | 235 | 11 | 20 | 63 | 309 | 39 | 60 |
| 1GUD_A–1RPJ_A | 355 | 4 | 12 | 92 | 265 | 24 | 80 |
| 1OTJ_D–1GY9_A | 264 | 6 | 4 | 63 | 303 | 33 | 30 |
| 1GQZ_A–2GKE_A | 292 | 7 | 20 | 66 | 263 | 41 | 55 |
| 1URP_D–2DRI_A | 398 | 13 | 9 | 48 | 184 | 37 | 51 |
| 1FSF_A–1FQO_B | 230 | 8 | 5 | 56 | 255 | 31 | 36 |
| 1G6W_D–1K0B_C | 146 | 1 | 1 | 29 | 175 | 27 | 16 |
| 1WD7_B–1WCW_A | 243 | 9 | 16 | 59 | 211 | 38 | 31 |
| 1AW2_A–1AW1_B | 303 | 3 | 4 | 92 | 170 | 28 | 33 |
| 1GQN_A–1QFE_B | 278 | 1 | 11 | 43 | 219 | 5 | 24 |
| 1YL5_B–1YL7_A | 253 | 5 | 6 | 56 | 198 | 24 | 12 |
| 1S2O_A–1TJ5_A | 165 | 4 | 8 | 48 | 276 | 24 | 54 |
| 1NJG_B–1NJF_A | 112 | 3 | 1 | 26 | 261 | 37 | 38 |
| 2LAO_A–1LAH_E | 209 | 2 | 6 | 35 | 205 | 20 | 51 |
| 1HW1_B–1H9G_A | 104 | 8 | 1 | 38 | 129 | 24 | 16 |
| 1AKZ_A–1SSP_E | 162 | 3 | 7 | 31 | 211 | 18 | 43 |
| 1ZOL_A–1O03_A | 131 | 0 | 8 | 27 | 204 | 28 | 57 |
| 2RCS_H–1AJ7_H | 185 | 7 | 10 | 62 | 247 | 41 | 35 |
| 1TJD_A–1EEJ_B | 116 | 3 | 5 | 33 | 188 | 37 | 18 |
| 4AKE_B–2ECK_B | 182 | 12 | 11 | 41 | 69 | 28 | 50 |
| 1MO7_A–1MO8_A | 149 | 30 | 20 | 29 | 112 | 75 | 74 |
| 1VIY_C–1VHL_A | 178 | 3 | 6 | 35 | 84 | 7 | 10 |
| 1LMZ_A–1P7M_A | 87 | 8 | 13 | 36 | 72 | 36 | 61 |

| | | | | | | | |
|---|---|---|---|---|---|---|---|
| **1PDB_A–1YHO_A** | 187 | 18 | 15 | 53 | 106 | 53 | 28 |
| **1EX6_B–1EX7_A** | 2 | 0 | 0 | 65 | 243 | 37 | 38 |
| **2IN2_A–2B0F_A** | 86 | 8 | 6 | 23 | 260 | 84 | 82 |
| **2LKC_A–2LKD_A** | 111 | 17 | 21 | 49 | 96 | 104 | 79 |
| **1IGP_A–2AU6_A** | 164 | 1 | 1 | 28 | 186 | 11 | 25 |
| **2LHS_A–2BEM_A** | 174 | 9 | 16 | 28 | 157 | 45 | 54 |
| **1F3Y_A–1JKN_A** | 107 | 9 | 3 | 12 | 180 | 62 | 67 |
| **2F63_A–1EQM_A** | 150 | 9 | 14 | 22 | 92 | 104 | 27 |
| **1TFU_A–3UC5_A** | 102 | 0 | 2 | 41 | 149 | 17 | 18 |
| **1XSA_A–1XSC_A** | 76 | 7 | 7 | 39 | 176 | 54 | 47 |
| **1IJA_A–2KID_A** | 148 | 7 | 23 | 15 | 84 | 58 | 47 |
| **1ORM_A–1QJ8_A** | 119 | 6 | 81 | 38 | 74 | 74 | 104 |
| **2KQ2_A–2KW4_A** | 32 | 45 | 38 | 94 | 16 | 81 | 170 |
| **1W4U_A–1UR6_A** | 72 | 12 | 7 | 22 | 80 | 115 | 63 |
| **1CFC_A–5DOW_A** | 46 | 8 | 12 | 4 | 12 | 54 | 40 |
| **1DMO_A–3CLN_A** | 49 | 14 | 12 | 4 | 11 | 55 | 22 |
| **2KXL_A–2K0G_A** | 125 | 17 | 16 | 8 | 40 | 47 | 59 |
| **1PFL_A–1FIL_A** | 93 | 9 | 6 | 15 | 117 | 61 | 29 |
| **1FMF_A–1ID8_A** | 87 | 52 | 16 | 27 | 32 | 106 | 85 |
| **2UZ5_A–2VCD_A** | 124 | 8 | 38 | 18 | 61 | 45 | 62 |
| **2P3M_A–2VBT_A** | 124 | 4 | 8 | 25 | 90 | 15 | 54 |
| **1JFJ_A–1JFK_A** | 47 | 4 | 5 | 15 | 30 | 14 | 7 |
| **1MX7_A–1MX8_A** | 129 | 12 | 17 | 10 | 35 | 30 | 28 |
| **2K43_A–2K8R_A** | 25 | 2 | 3 | 7 | 114 | 133 | 107 |
| **1AEL_A–1URE_A** | 130 | 12 | 19 | 12 | 52 | 82 | 36 |
| **1I56_A–1EL1_A** | 13 | 3 | 0 | 2 | 133 | 64 | 35 |
| **1MUT_A–1PUN_A** | 72 | 15 | 23 | 26 | 78 | 75 | 84 |
| **2JU3_A–2JU8_A** | 116 | 22 | 5 | 16 | 22 | 47 | 24 |
| **1EAL_A–1EIO_A** | 124 | 4 | 17 | 10 | 32 | 31 | 33 |
| **1O1U_A–1O1V_A** | 129 | 6 | 6 | 14 | 31 | 32 | 37 |
| **2L68_A–2LKK_A** | 112 | 5 | 11 | 21 | 51 | 14 | 26 |
| **2AI6_A–2OZW_A** | 106 | 7 | 6 | 22 | 124 | 42 | 45 |
| **1NTR_A–1KRX_A** | 97 | 11 | 28 | 19 | 48 | 66 | 97 |
| **2D9E_A–2RS9_B** | 57 | 3 | 5 | 14 | 38 | 13 | 9 |
| **1JM4_B–1WUM_A** | 58 | 3 | 3 | 18 | 50 | 25 | 23 |
| **2JWW_A–1RTP_1** | 24 | 3 | 4 | 7 | 59 | 75 | 29 |
| **2NLN_A–1RRO_A** | 25 | 3 | 2 | 4 | 52 | 65 | 35 |
| **2L50_A–2L51_B** | 11 | 8 | 5 | 20 | 15 | 14 | 25 |
| **1HSI_B–1HSH_D** | 24 | 0 | 1 | 14 | 131 | 12 | 7 |
| **2CJO_A–1ROE_A** | 63 | 26 | 8 | 15 | 49 | 73 | 90 |
| **1C54_A–1RGH_B** | 61 | 1 | 3 | 11 | 71 | 27 | 10 |
| **1K2H_A–1ZFS_B** | 13 | 5 | 14 | 13 | 12 | 70 | 15 |
| **1SYM_B–1XYD_B** | 13 | 6 | 13 | 10 | 12 | 62 | 12 |
| **1BV2_A–1UVB_A** | 21 | 1 | 2 | 6 | 69 | 36 | 23 |
| **1LIP_A–1JTB_A** | 13 | 1 | 7 | 8 | 57 | 69 | 43 |
| **2FHM_A–2HLT_A** | 117 | 4 | 16 | 16 | 38 | 22 | 25 |
| **1SKT_A–1TNQ_A** | 20 | 12 | 8 | 8 | 13 | 55 | 13 |
| **1GH1_A–1CZ2_A** | 19 | 2 | 5 | 6 | 46 | 59 | 37 |
| **2CG7_A–2CG6_A** | 17 | 0 | 0 | 10 | 85 | 14 | 12 |

### Supplementary Figures

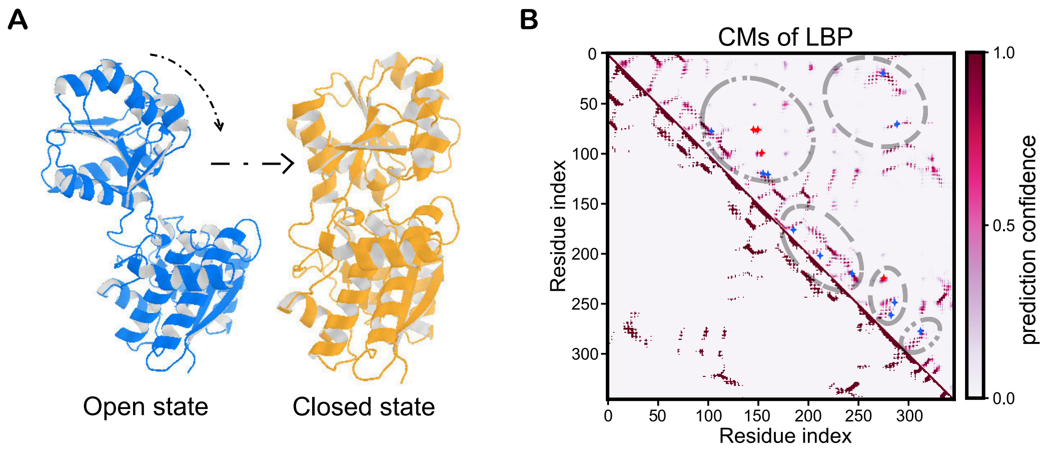


**Figure S1:** More details on the analysis of leucine-binding protein (LBP). (A) Apo and holo structures of LBP showed a clear open–closed transition. (B) Comparison between the contact map from the holo state (TCM1, lower left) and the predicted contact map from its sequence (PCM, upper right) shows additional contacts of high prediction confidence (gray circle), which may represent new contacts in the alternative conformation. The star points are in correspondence with the residue pairs in Table 2, where red stars represent the selected residue pairs in bold and blue ones for the rest.

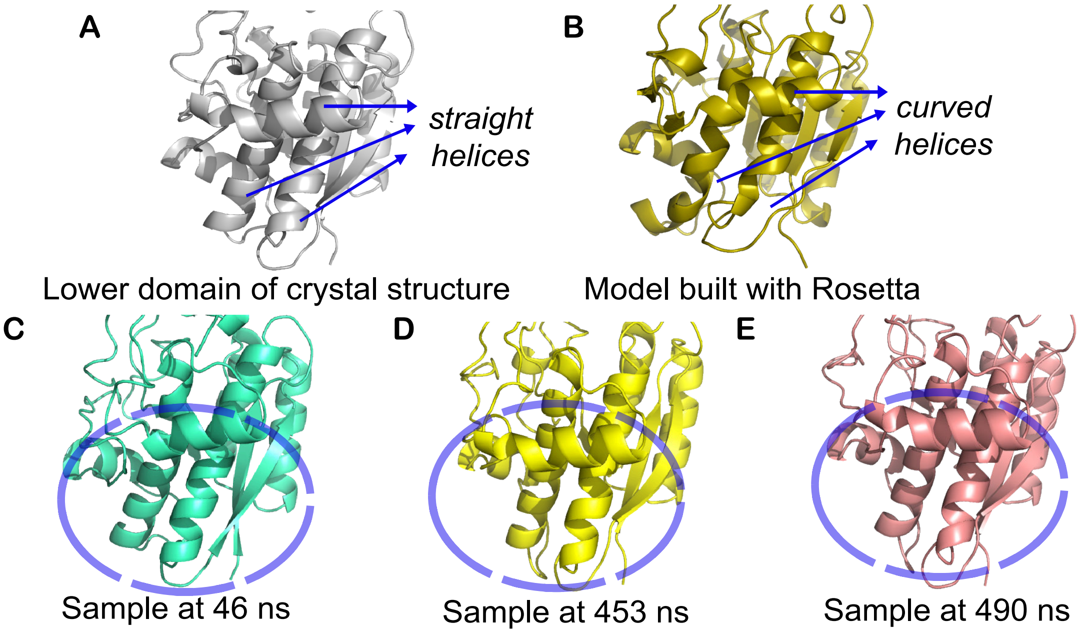


**Figure S2:** Structural refinement of the predicted closed state of LBP in the MD simulations. One of the main improvements lied on the helices of the lower domain, which were straight in the crystal structure (A) but curved in the Rosetta-built model (B). As shown in (C)–(E), the MD-sampled conformations showed clear refinement for these helices.
